# Disentangling the relative importance of spatio-temporal parameters and host selectivity in shaping AMF communities in temperate forests

**DOI:** 10.1101/2021.02.16.431389

**Authors:** Leonie Grünfeld, Magkdi Mola, Monika Wulf, Stefan Hempel, Stavros D Veresoglou

## Abstract

Many woody and herbaceous plants in temperate forests cannot establish and survive in the absence of mycorrhizal associations. Here, we address and hierarchize how Glomeromycota, a group of soil-borne fungi, forming a ubiquitous type of mycorrhiza, varied in a temperate forest patch in Germany with time, space, plant hosts but also the proximity to Glomeromycota-associating woody species. The communities of Glomeromycota in our study were non-random. We observed that space had a greater impact on fungal community structure than either time or proximity to Glomeromycota-associating trees but unlike host identity did not alter Glomeromycotan richness. The set of parameters which we addressed has rarely been studied together and we believe that the resulting ranking could ease prioritizing some of them to include in future surveys. Glomeromycota are crucial for the establishment of understory plants in temperate forests making it desirable to further explore how they vary in time and space.

## Introduction

Arbuscular mycorrhizal fungi (AMF) are globally distributed symbiotic fungi, which at large spatial scales show non-random distribution patterns, explained mainly by abiotic predictors such as pH ([1,2]), soil properties [3] and climatic conditions [4] but also host selectivity [5]. Most studies addressing AMF, however, can only explain a small fraction of AMF community variance, suggesting that AMF communities are subject to a high degree of stochasticity (i.e. the fraction of community variance not explained by deterministic processes; Appendix I; [6,7,8]). Assaying stochastic drivers in large-scale mycorrhizal studies is tricky, because they can inflate the sequencing effort required. We aimed here at ranking the relative importance, in terms of shaping AMF communities, of a set of stochastic and biotic drivers rarely assayed together so as to smooth the way of integrating stochastic predictors into mainstream studies on mycorrhiza.

We present a spatio-temporal study in a forest where we address the relative importance of (i) physical distance, (ii) sampling time, (iii) host selectivity and (iv) proximity to AMF-associating woody species [9,10] in shaping the AMF community structure of the understory. To the best of our understanding, no other mycorrhizal study to date has simultaneously studied this specific set of parameters, even though studying subsets of them have generated highly-valued expectations. Davison et al. [11], for example, studied the effects of seasonality and spatial structure in an Estonian temperate forest to observe considerable spatial heterogeneity in AMF species distribution, but minimal changes over the duration of a growth season. Dumbrell et al. [4], by contrast, observed pronounced temporal changes in the composition of AMF grassland root communities over a single growth season. Su et al. [12] addressed the relative strength of host selectivity and seasonality to show that in the particular system of Inner Mongolia, seasonality masked any host preferences across five hosts. As a result, the first expectation of our study was that physical distance would exceed in relative importance temporal variance in shaping AMF communities (*Hypothesis One*; [11]). It is likely that woody plants in those earlier studies had such strong effects on the understory because they had acted as islands of AMF propagules [10]. If it is the presence of AMF-associating woody species which mainly shapes the regional pool of AMF species, then we might expect a lower, compared to grasslands, relative importance of host selectivity across the understory plants. We thus additionally hypothesized that proximity to an AMF-associating woody species would alter AMF community structure more than host selectivity does (*Hypothesis Two*). We addressed these two hypotheses in a 625 m^2^ forest site in the Elbe-Weser region in North-West Germany which we monitored over two years, totalling four harvests of root material.

## Materials and Methods

### Study site

The study site is a floristically well described [13,14] temperate European deciduous forest in northwest Germany (53.44°N; 9.49°E). Biophysical characteristics were as follows: Soil pH was between 4.6 and 5.6, P between 9 and 28 mg kg-1, K between 36 and 112 mg kg-1, Ca between 1396 and 1885 mg kg-1 and Mg between 78 and 98 mg kg-1 (Table S2). We divided the site into 25 5 x 5 m rectangular plots, assessed in situ woody vegetation cover of AMF-associating plants per plot and classified them accordingly into *high* and *low* (≥ 15 % and < 15 % AMF-associating woody cover, respectively; Fig. S1, Table S1; we rationalize the choice of the threshold in Fig. S2) classes. Based on earlier observations in the area, understory plants associating with AMF occurred at higher frequency [9] and were colonized more extensively by AMF [10] in the proximity of woody plants forming arbuscular mycorrhiza.

### Sampling Design

In May and September of 2017 and 2018, we collected *Hedera helix* L. (from now on *Hedera*) roots from pairs of *high* and *low* plots (78 samples in total). In September 2018, we additionally collected roots of *Euonymus europaeus* L. (19 samples in total; from now on *Euonymus*). Roots were excavated up to a depth of about 10 cm from two plant individuals per plot, which were processed independently. The root samples were subsequently cleaned with water and transferred into a falcon tube with 95 % ethanol. In total, there were 97 root samples from two plant species. In specific we collected 12 root samples from 6 plots in May 2017, 19 samples from 9 plots in September 2017 and May 2018 – one plot was represented with a single sample in each case because there were no other plants occurring at a distance higher than 50 cm at the plot. There were 26 *Hedera* root samples from 14 plots (in two cases there was a single sample per plot) and 17 *Euonymus* samples from 10 plots (in three cases there was a single sample per plot) in September 2018.

### Molecular analyses and bioinformatics

Roots were transported to the lab on ethanol at 4°C and stored at −20°C. Root samples were freeze-dried and homogenized with a Retsch Mixer Mill MM 400 using metal balls of 1 mm diameter. DNA was extracted from 30 mg ground root material per sample with the DNeasy® PowerPlant® Pro Kit (Qiagen, Venlo, the Netherlands) and amplified with the Glomeromycota-18S-rRNA gene targeting primer pair NS31-AML2 [15] extended with the adaptors p5 (NS31) and p7 (AML2; [16]). The amplification conditions were as follows: each of the 25 µl PCR reactions contained 1 µl DNA template, 2.5 µl (0.3 µM of each primer) primer mix, 0.25 µl KAPA HiFi DNA polymerase (1 U/µl), 0.5 µl KAPA dNTP mix (10 µM), 5 µl 5X KAPA HiFi Fidelity Buffer and 15.75 µl nuclease-free water. The PCR reactions were performed with a Biometra-Ton thermal cycler (Analytik Jena, Jena, Germany) under the following conditions: Initial denaturation at 95 °C for 2 min, 35 cycles with a denaturation phase of 98 °C for 45 s, an annealing phase of 65 °C for 45 s and an extension phase of 72 °C for 45 s and final elongation at 72 °C for 10 min. Samples that did not perform well on this initial PCR (∼40% of the samples) were amplified instead with a GC-rich buffer from the kit. Four out of 97 samples did not show bands during gel electrophoresis and were excluded from further analysis. The NS31-AML2 amplicons were purified with the NucleoSpin® gel and PCR clean-up kit (Macherey-Nagel, Düren, Germany). For indexing purposes we used Miseq specific adaptors (NuGen) which we ligated to our products with an additional PCR. The PCR master mix for indexing consisted of 1 µl of the purified PCR template, 2.4 µl of the primer mix, 0.25 µl Phusion® high-fidelity DNA polymerase (BioLabs), 0.5 µl dNTPs (10 µM), 5 µl 5X Phusion® HF buffer and 15.85 µl nuclease-free water per 25 µl reaction. After indexing PCR – thermocycling settings: 95 °C for 3 min, 15 cycles of 98 °C for 30 s, 55 °C for 30 s and 72 °C for 30 s, and 72 °C for 5 min – the DNA fragments were separated by gel electrophoresis to check the signal strengths. We used MiSeq Illumina chemistry to sequence the amplicons. We processed the libraries with the uPARSE pipeline [17] with uSearch v 10.0.240. In brief, forward and backward reads were merged with the fastq_mergepairs command, primers were stripped and sequence pairs with a length shorter than 400 bp or more than 1 expected error were filtered out. We used the cluster_otus command to construct the OTU table. Representative OTUs sequences were blasted against MaarjAM [18] and non-specific to Glomeromycotina OTUs (i.e. < 97.5 % similarity or < 99 % coverage) were excluded from further analyses. We then rarefied to 2350 reads per sample, which excluded 2 samples from further analysis (i.e. analysis was carried out to the remaining 91 samples).

## Statistical analyses

### Null model analysis - to what degree AMF distributions were random?

To address the degree to which AMF communities were random, we engaged into a null model analysis with the R package EcoSimR [19]. We compared C score (i.e. checkerboards) occurrences in our dataset in comparison (i.e.we z-score standardized effect sizes (SES) in relation to the set of simulated community matrices) to distributions of 1000 random matrices that were generated with the *sim*4 algorithm. We used presence-absence data and kept in the community table the total number of row sums fixed, describing how often species occurred, and proportional to those observed the column sums, describing differences across samples. The *sim*4 algorithm effectively controls for Type I and II statistical errors and has been proposed for scenarios where some rare species have occasionally been scored as absent even though present (i.e. incomplete lists; [20]). Negative standardized effect sizes below −1.96 manifest aggregation of species within samples/sites, positive values above 1.96 segregation of species within samples, whereas values between −1.96 and 1.96 a random species distribution. Pooling together heterogeneous samples might bias the results towards appearing less random [21]. We thus additionally assessed null model statistics for several subsets of the combined community matrix.

### Hypothesis One: physical distance is more important than temporal variance in structuring AMF

To address this hypothesis we (i) visualized the raw data; (ii) calculated effect sizes for the major drivers of AMF community structure; and (iii) presented as a key result a summary for some characteristic groups of samples of community structure information at the family level. First, we carried out a Principal Components Analysis (PCA) to visualize clustering patterns across the samples. Second we presented how effect sizes differed with our variables of interest. To avoid statistical pitfalls from partitioning variance in spatio-temporal designs (e.g. [22]; Appendix II), we visualized relative effect sizes by means of Bray-Curtis distances and only supplementary fitted a predictive model in the form of a redundancy analysis (RDA). To compare effect sizes, we randomly paired samples sharing specific attributes 9999 times and quantified Bray-Curtis distances. Third, we summarized how AMF community structure differed with each of the predictors, by generating bar plots with relative abundance information on each AMF family. We finally created a heatmap presenting the frequencies with which individual AMF taxa were observed in habitats of specific attributes.

### Hypothesis Two: proximity to an AMF-associating woody species would alter AMF community structure more than host selectivity does

We first carried out a repeated-measures ANOVA to compare diversity metrics (i.e. richness, Shannon diversity and Pielou evenness) between the two types of habitats, which we fitted as a mixed-effects model. The response variable was the diversity metric; host species and *low* vs. *high* type of habitat were the predictors and time was the repeated measures parameter. We additionally used in our repeated-measures ANOVA spatial autocorrelation parameters to correct for spatial dependencies. To address whether the communities in these two habitats differed in relation to how aggregated/segregated they were, we further compared the respective SES which we yielded from our null model analysis. We created a venn diagram depicting how host selectivity and proximity to AMF-associating woody species influence AMF community structure to visualize compositional differences. We finally carried out an indicator species analysis (we used the package indicspecies in R; [23]) in relation to the following classes: the two plant species, the two habitat types (i.e. *high* vs *low*) and their meaningful combinations.

## Results

### Overall Statistics

Out of 853,811 quality controlled reads, 696,451 described 32 Glomeromycota-specific OTUs (Table S3). Eighteen of them belonged to Glomeraceae, six to Claroideoglomeraceae, five to Archaeosporaceae, two to Diversisporaceae and one each to Gigasporaceae and Acaulosporaceae. We rarefied sequencing depth to 2350 reads which resulted in the exclusion of two samples. AMF richness varied between 4 and 17 OTUs per sample (median: 10 OTUs with the quartiles being 8 and 12; Fig. S3). Richness only differed with plant species (*t* = −4.44, *P <* 0.0001; when we narrowed observations to those from the forth harvest the respective statistics were *t* = −3.14, *P =* 0.003; Fig. S3): *Euonymus* plants contained on average 8.2 AMF taxa, whereas *Hedera* plants 10.54.

The indicator species analysis classified 5 out of the 55 species as indicators. OTU2 (Glomeraceae; *P*=0.045) was an indicator of *Euonymus* communities and OTU70 (Glomeraceae, *P*<0.001) an indicator of *Euonymus* community at *low* plots. OTU8 (Claroideoglomaceae; *P*=0.001) and OTU13 (Acaulosporaceae, *P*<0.01) were indicators of *Hedera* communities whereas OTU19 (Diversisporaceae; *P*=0.038) specifically associated with *Hedera* at *high* plots.

### Null model analysis - to what degree AMF distributions were random?

In all our tests we observed a significant species aggregation (Fig. 1). The standardized effect sizes (SES) varied between −10.90 (combined community matrix) and −2.4 (*Hedera* roots in May 2017). *Hedera* roots from *low* plots (SES = −8.19) showed more aggregated AMF communities than their representatives from *high* plots (SES = −4.81; any differences in the statistics exceeding 1.96 are significant). Also AMF communities in *Hedera* were more aggregated in autumn, compared to spring (the mean SES statistic for spring was −2.98 whereas for autumn it was −5.25). The results in SES statistics could not be explained based on sampling intensity (i.e. number of individuals assayed).

**Fig. 1.**
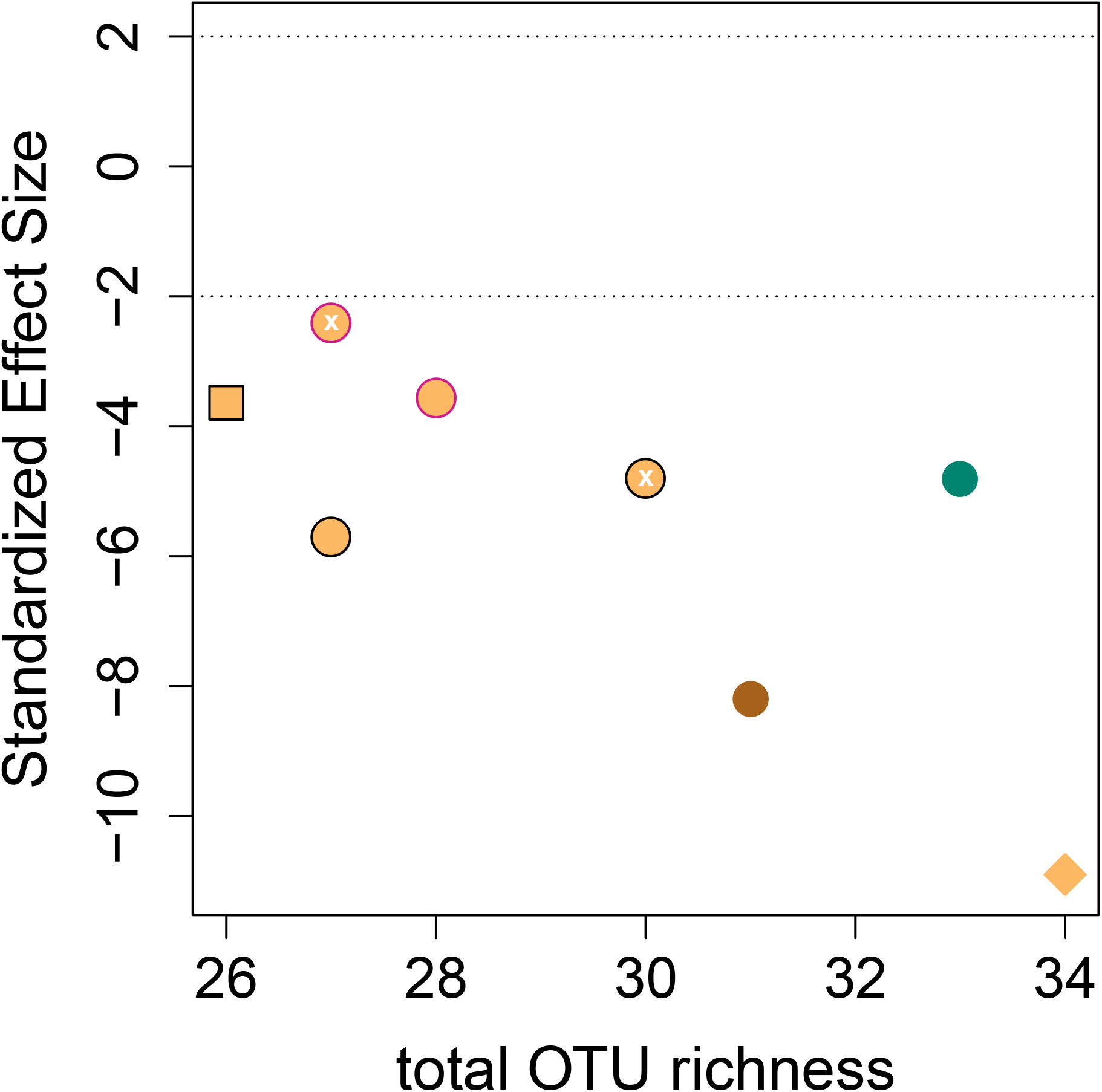
Standardized effect sizes of observed checkerboard scores compared against null models which were generated with the sim4 algorithm (*y*-axis). We plotted these values against total OTU richness of the respective subsets of the dataset. The two discontinuous lines highlight confidence intervals within which the community matrix can be considered random. The green point represents samples from high plots, the brown from low plots and the orange points from combinations of the two. A pink border was used for spring and a black for autumn; we used no border in cases where we pooled samples from spring and autumn. We used white “x” symbols to highlight the location in the panel of samples taken over the first year. The square describes samples on *Euonymus* whereas circles those on *Hedera*. The diamond shows the complete data set. Differences in standardized effect sizes above 1.96 are significant at a 0.05% confidence level.

### Hypothesis One: physical distance is more important than temporal variance in structuring AMF

Our Principal Components Analysis on Hellinger-transformed occurrence data hinted that any differences in AMF community structure across the samples were subtle (Fig. 2a). The resulting histograms overlapped considerably but spatial structure induced stronger effect sizes than temporal variability. In addition, community changes within a growth season were subtle (Fig. 2b). We also observed that the two plant species (Fig. 2b) shared more similar communities than expected by chance and that it was low plots that had the most divergent AMF communities. (Fig 2c). *Euonymus*-associated AMF communities were dominated by Glomeraceae (94.8 % on average compared to a maximum of 86 % in *Hedera*; Fig. 2c). High occurrence of Glomeraceae was also observed at low AM plots (averaging 85.5 %). Relative abundance differences of families were considerably more pronounced across years than across seasons (Fig.2c). In the redundancy analysis with the drivers as predictors, we found that year, plant species and spatial autocorrelation axes explained AMF community shifts whereas season had no effect. AM-plant cover shared considerable variance with other predictors and significance depended on the ranking with which it was included in the predictors (Appendix III).

**Fig. 2.**
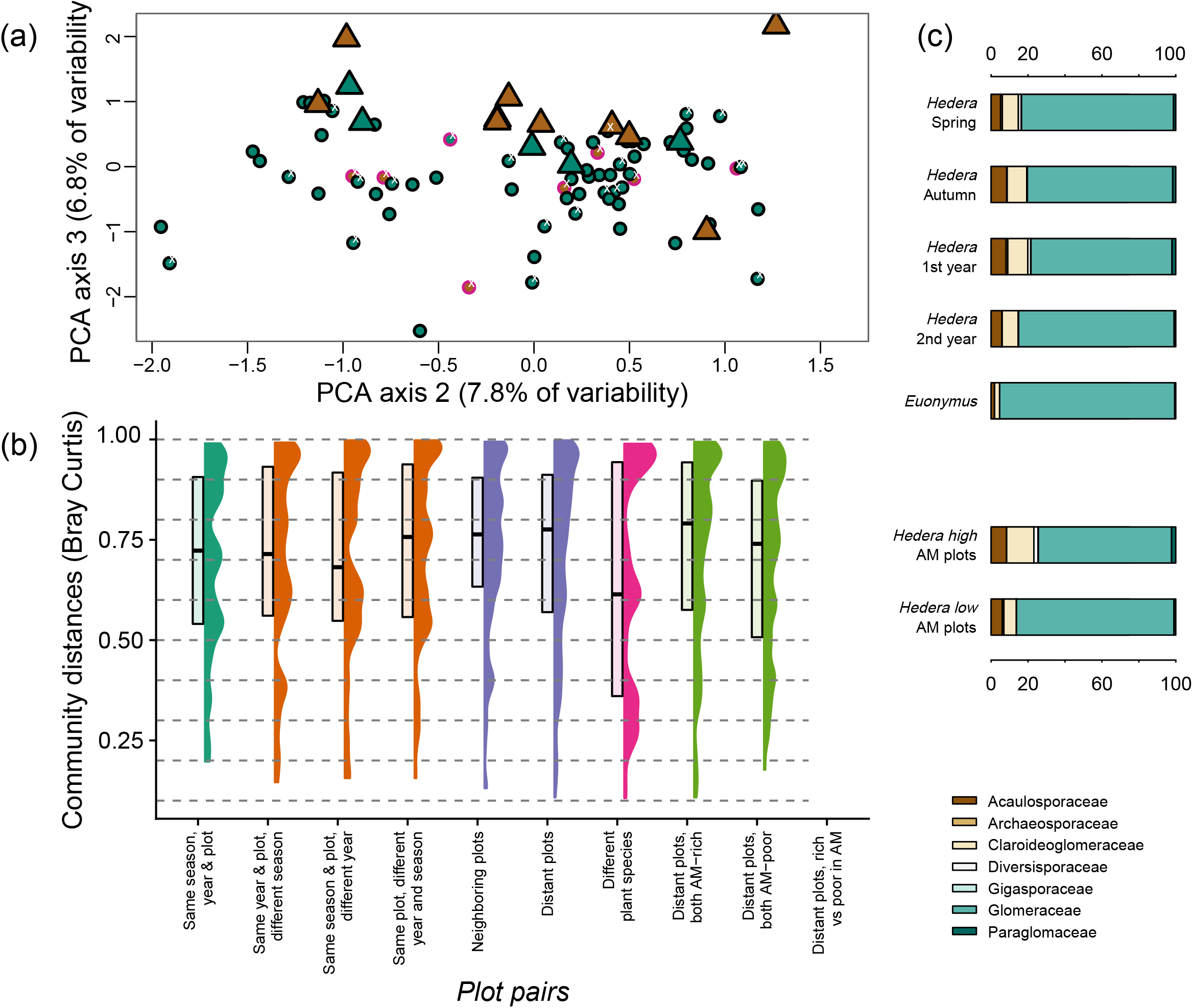
(a) Principal component analysis of Hellinger transformed AMF community data (we plotted respective diagrams with axis one, explaining 13.5 % of variability in Fig. S4 and Fig. S5). Green symbols represent samples from high plots whereas brown from low plots. A pink border was used for spring and a black for autumn. We used white “x” symbols to highlight the location in the panel of samples taken over the first year. Triangles describe samples on *Euonymus* whereas circles those on *Hedera*. (b) Distributions of pairwise community distances (Bray-Curtis distances) for a range of pairwise combinations (dark green: within plots sampled at the same time; orange: same plot differing in sampling time; purple: same harvest but different plot; pink: same plot in the 4^th^ harvest but different plant species; light green: same harvest but different plot grouped based on the relative coverage of AMF-associating woody plants.). Note that Bray-Curtis community distances between *Hedera* and *Euonymus* (in pink; same plot) were smaller than respective distances between individuals of *Hedera* (dark green) (c) Mean relative abundances of the seven AMF families (Acaulosporaceae, Archaeosporaceae, Claroideoglomeraceae, Diversisporaceae, Gigasporaceae, Glomeraceae, Paraglomaceae) grouped based on (up) the time of sampling and plant host and (bottom) our classification into high and low plots.

### Hypothesis Two: proximity to an AMF-associating woody species would alter AMF community structure more than host selectivity does

We observed no diversity differences in relation to proximity to AMF-associating woody species (Fig. 1; F_1,81_ = 0.048, *P* = 0.83; The only significant effect was that of plant species; F_1,81_ = 11.8, *P* < 0.001). Roots from *low* plots contained consistently more aggregated AMF communities than their representatives from *high* plots (Fig. 1). There were minor compositional distances between *high* and *low* plots with 7 OTUs being specific to high plots and 2 to low plots (Fig. 3a) We observed, by contrast, twelve OTUs to be specific to *Hedera* samples (Fig. 3a), which might have been because of the most extensive sampling of *Hedera* individuals. Observation frequency, for most taxa, was higher at *high* plots than at *low* plots (Fig. 3b).

**Fig. 3.**
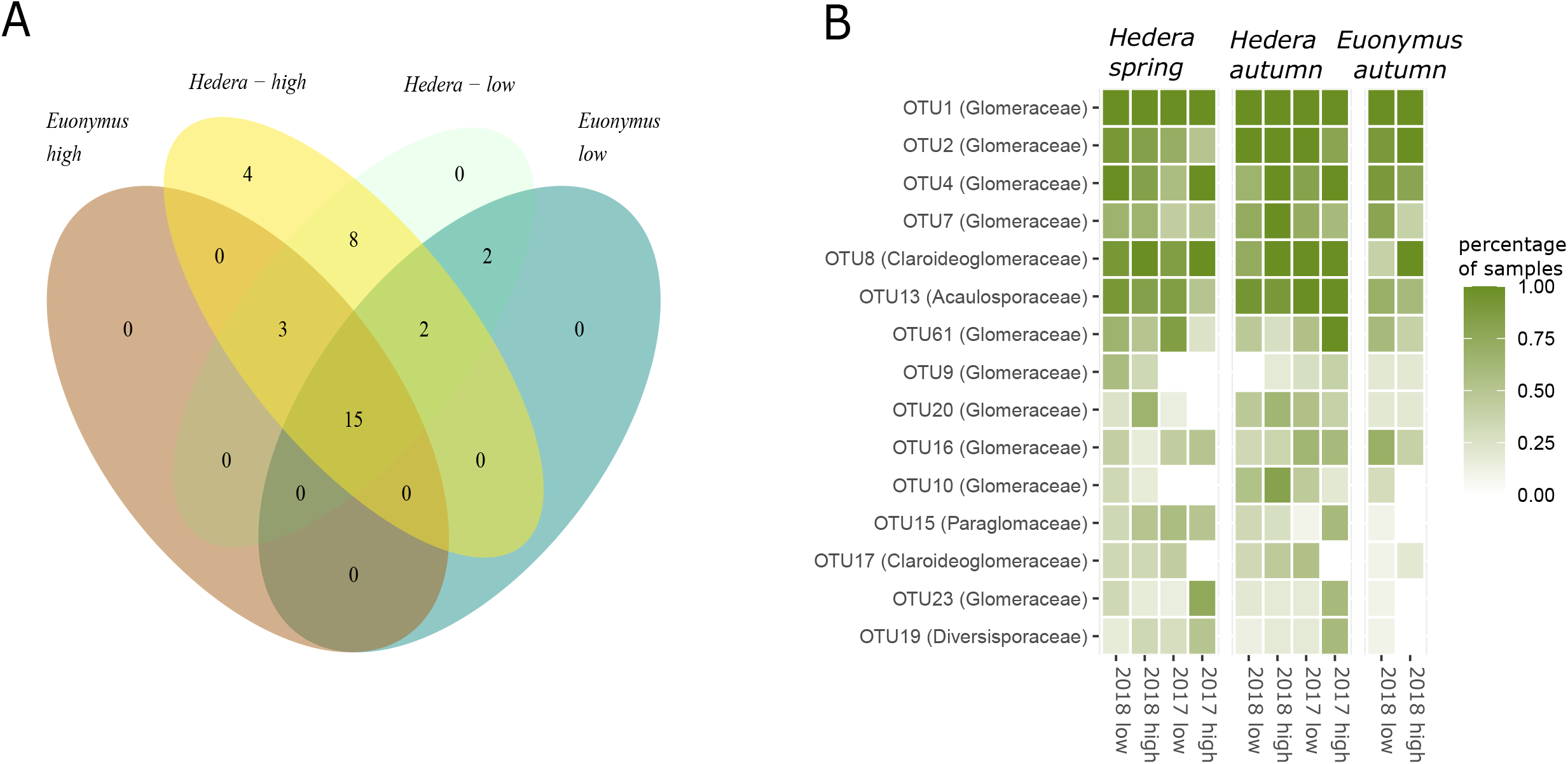
(A) Venn diagram depicting the distribution of OTUs across (i) *high* and *low* plots and (ii) the two plant hosts. Fifteen out of the thirty-two OTUs were observed in all four types of habitats. (B) Frequency of occurrence of the fifteen most abundant OTUs across ten groups of samples describing plant host, plot quality in relation to AMF abundance and season of sampling.

## Discussion

A take home message of our study is that, in agreement with *Hypothesis One*, physical distance in temperate forests exerts a stronger influence on AMF communities than either sampling time or host selectivity. We also show that temporal variability occurs across years and not across seasons. Hence, our data agree with Davison et al. [11] that there is low seasonality in forests in relation to AMF communities. The order of establishment of plant hosts, known as priority effects, could thus play an important role in structuring AMF communities [24]. In natural systems this most likely occurs in the beginning of the growing season. Even though this idea remains underexplored, it could potentially explain why the effect sizes for different years were larger than for different seasons.

Through our null model analysis, we deduced that the plant colonization patterns in our study had been non-random (even though AMF community differences with time, space and hosts were weak - Fig. 2a) and showed extensive aggregation of species, meaning that the OTUs co-occurred more often than expected by chance. That our null model analysis supported that AMF root community structure was not random, was not surprising (e.g. [25]). The outcome of co-occurrence analyses, however, depends strongly on how heterogenous the compared communities have been (but also on sampling intensity): relatively homogenous communities such as those in our study are more likely to show aggregation whereas heterogeneous pools of samples such as those analyzed with a comparable approach in Hu et al. [26] are more likely to show segregation. It was important in our study to first show that the community matrix at the used spatial scale had been non-random (and thus our study had enough resolution to address community variance patterns in AMF communities), before addressing how spatio-temporal parameters and host specificity explained the community variance. Additionally, through our null model analyses we could observe some overarching patterns such as that *low* plots hosted more aggregated AMF communities than *high* plots. Species aggregation patterns often suggest shared habitat requirements across species (compared to mechanisms such as competition and dispersal limitation which induce segregation e.g. [27]) which might have been stronger across AMF colonizing *Hedera* roots in *low* plots.

In our RDAs, we observed pronounced plant host effects on AMF diversity (Fig. S1), AMF community aggregation (Fig. 1) and community structure (Fig. 2). The present study obviously did not fully address the role of host selectivity: we only assayed two host plants and because of the low abundance of *Euonymus*, we only assayed respective individuals in the last harvest. This mainly served the purpose of showing the degree to which our observations with *Hedera* were generalizable. We believe, nevertheless, that we could still get a reasonable (and hopefully representative) picture of how host selectivity influences AMF communities. We present evidence, for example, that host selectivity has a strong influence on AMF richness (i.e. host identity was the only parameter in our analyses that had an effect on AMF richness). We found of special interest, though, that pairwise differences between species (*Hedera*– vs. *Euonymus*–associating AMF communities; Fig 2b, pink bar) were smaller than respective pairwise differences of conspecific individuals (randomly paired in rda models). There is evidence that phylogenetically divergent co-occurring plant species share more similar AMF communities than closely related species [28] and our analysis hints towards this direction. Remarkably, most larger-scale studies found no evidence for host selectivity (e.g. [29]). This could mean that abiotic conditions mask host selectivity at larger scales. Alternatively, inconspicuous factors at a smaller scale driven by the environment such as priority effects or the availability of AMF propagules could modify how plant species select for AMF communities.

Contrary to our expectations that *low* and *high* plots would host distinct AMF communities (*Hypothesis Two*), we only observed small associated differences in diversity and the factor AM plant cover in the RDA was only conditionally significant (Fig. 2a; Appendix III). This was despite the fact that AMF communities across low plots appeared more divergent (Fig. 2b; light green middle bar) and that we observed differences in relation to the aggregation patterns (Fig. 1). In Grünfeld et al. [10] we had observed pronounced differences in root colonization between *high* and *low* plots across forests in the specific area but we had worked at a relatively larger spatial scale. AMF can grow vegetatively to distances of about 50 cm [30]but they could also potentially disperse by other means such as air and animal vectors [31]. We may have thus missed the relevant spatial scale, or differences in relation to the mycorrhizal state of the canopy impact percentage colonization to a larger degree than AMF community structure.

We compare and rank relative effect sizes of drivers of AMF community structure operating at a small spatial scale (as compared to soil properties and climatic variables that operate at larger scales) that have rarely been addressed simultaneously. Several authors such as Dumbrell et al. [6] have highlighted the need to better understand stochastic processes in AMF and our study presents a ranking exercise which contributes towards this direction.

## Acknowledgements

The authors acknowledge funding from the Deutsche Forschungsgemeinschaft Project Metacorrhiza (VE 736/2-1) awarded to SDV. We thank Matthias Rillig for comments to an earlier version of the manuscript.

## Author Contributions

Conceived the study, did the bioinformatics and statistical analyses: SDV; Carried out the harvests: LG, MW, SDV; Did the molecular analyses LG, MM, Wrote the paper LG with contributions from SDV; All authors commented on the manuscript and approved the final version.

