## Appendix for "Disentangling the relative importance of spatio-temporal parameters and host selectivity in shaping AMF communities in temperate forests"

**Fig. S1** Schematic plan of the study site

**Fig. S2** Choosing a AM-coverage threshold

**Fig. S3** Richness statistics from all plots and harvests

**Fig. S4** PCA of AMF community data (first and second axes)

**Fig. S5** PCA of AMF community data after excluding outliers

**Table S1** AM plant cover and sampling information

**Table S2** Soil properties of the sampling site

**Table S3** OTU ID information

**Appendix I** Defining stochasticity

**Appendix II** Limitations of existing multivariate techniques in addressing spatio-temporal designs

**Appendix III** Statistical analyses

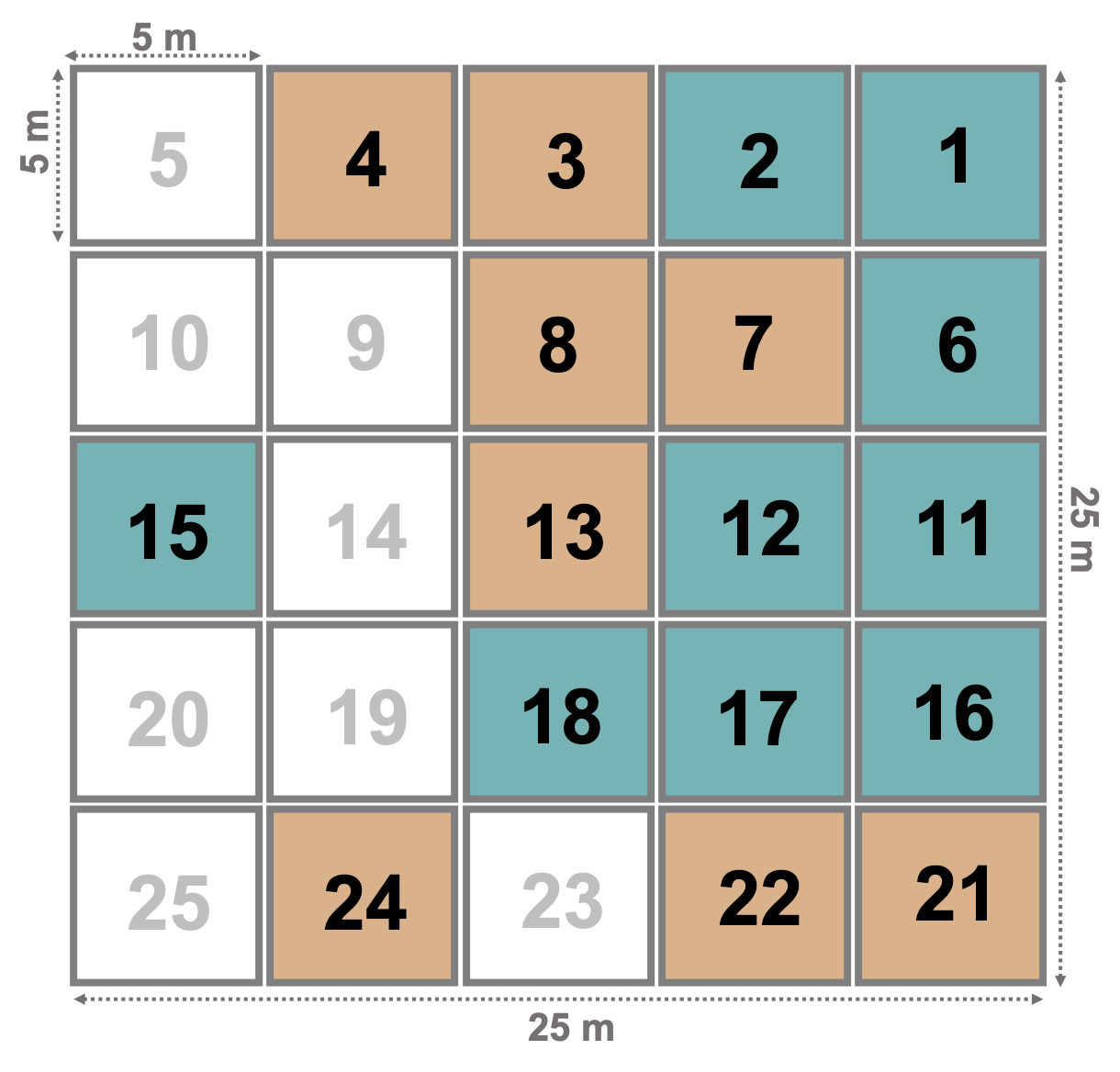

**Fig. S1** Schematic plan of the 25 5 m x 5 m plots within the study site. Green squares indicate plots with *high* (≥ 15 % AM plant cover) and brown squares indicate plots with *low* (< 15 % AM plant cover). White squares indicate plots where no root samples were collected.

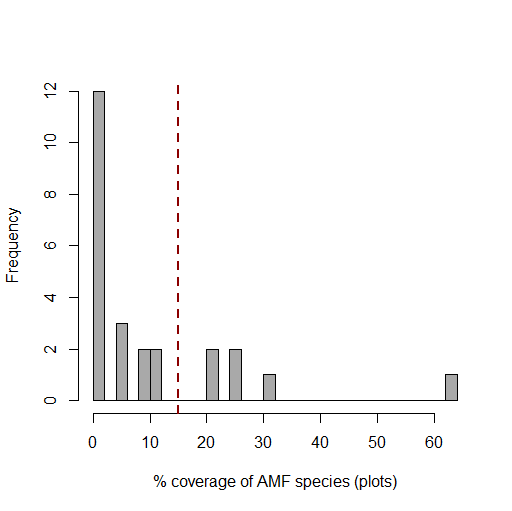

**Fig. S2** Histogram depicting the classification threshold of 15% in relation to AM-host coverage across plots. Plots with over 15% AM-host coverage were classified as *high* plots whereas those with a lower % AM-host coverage as *low*.

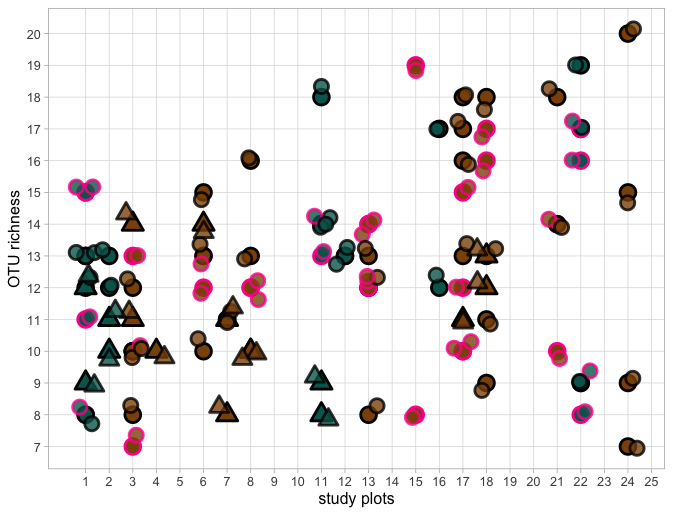

**Fig. S3** Richness statistics based on numbers of OTUs from all plots and harvests. Green symbols represent samples from high plots whereas brown from low plots. A pink border was used for spring and a black for autumn. Triangles describe samples on *Euonymus* whereas circles those on *Hedera*.

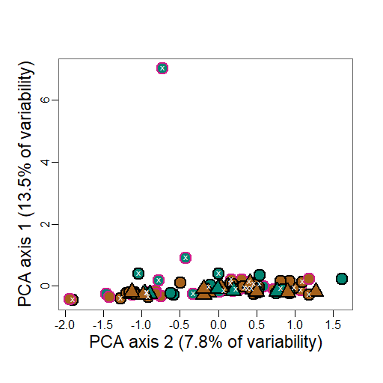

**Fig. S4** Principal component analysis of Hellinger transformed AMF community data with axis one explaining 13.5 % of variability. Green points represent samples from high plots whereas brown from low plots. A pink border was used for spring and a black for autumn. We used white “x” symbols to highlight the location in the panel of samples taken over the first year. Triangles describe samples on *Euonymus* whereas circles those on *Hedera*. In Fig. 2a we display axes two and three because after excluding the outlier in axis one the rescaled variances for the three first axes are 1.03% - axis 1, 7.84% - axis 2 and 6.88%.

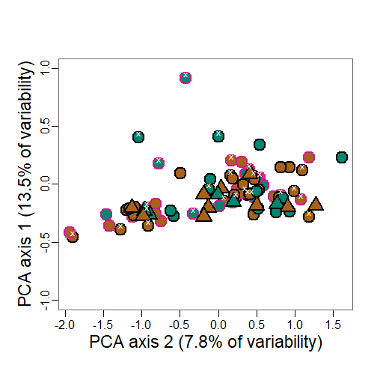

**Fig. S5** Principal component analysis of Hellinger transformed AMF community data with axis one explaining 13.5 % of variability (see Fig. S4) after the exclusion of outliers. Green points represent samples from high plots whereas brown from low plots. A pink border was used for spring and a black for autumn. We used white “x” symbols to highlight the location in the panel of samples taken over the first year. Triangles describe samples on *Euonymus* whereas circles those on *Hedera*. In Fig. 2a we display axes two and three because after excluding the outlier in axis one the rescaled variances for the three first axes are 1.03% - axis 1, 7.84% - axis 2 and 6.88%.

**Table S1** Visually assessed AMF-associating plant cover across the 25 plots presented in Fig. S1. We used a % cover threshold of 15% to classify them into *high* and *low* plots (see Fig. S2). Roots were collected from 17 out of the 25 plots. The four harvests took place on 31^st^ of May 2017, 27^th^ of September 2017, 31^st^ of May 2018 and 25^th^ of September 2018. We progressively made the harvests most inclusive by sampling plots where the two species had a high relative abundance. This was done to avoid issues arising from the two plants getting locally extinct.

| plot within site | % cover  of AM-associating woody plants | classification  AM plant cover | harvests where  *Hedera helix* was sampled | sampling  *Euonymus europaea*  In 4th harvest |
| --- | --- | --- | --- | --- |
| 1 | 26 | high | 1-4 | yes |
| 2 | 31 | high | 4 | yes |
| 3 | 10 | low | 1-4 | yes |
| 4 | 1 | low | NA | yes |
| 6 | 6 | low | 2-4 | yes |
| 7 | 11 | low | 4 | yes |
| 8 | 5 | low | 2-3 | yes |
| 11 | 22 | high | 2-4 | yes |
| 12 | 21 | high | 4 | no |
| 13 | 12 | low | 1-4 | no |
| 15 | 1 | low | 1 | no |
| 16 | 64 | high | 4 | no |
| 17 | 1 | low | 1-4 | yes |
| 18 | 2 | low | 2-4 | yes |
| 21 | 1 | low | 3-4 | no |
| 22 | 26 | high | 1-4 | no |
| 24 | 2 | low | 2,4 | no |

**Table S2** Soil pH (measured in 0.01m CaCl_2_) and nutrient levels of phosphorus, potassium, calcium and magnesium at the sampling site. Measurements were taken at three different soil depths by Wulf [1]. The soil constitutes of a humid to waterlogged pseudogley [2]. Since we worked at a relatively small spatial scale we expected soil characteristics to be consistent across plots.

| soil depth | pH | P [mg/100g] | K [mg/100g] | Ca [mg/100g] | Mg [mg/100g] |
| --- | --- | --- | --- | --- | --- |
| 0 - 5 cm | 4.6 | 2.8 | 11.2 | 139.6 | 9.8 |
| 5 - 10 cm | 5.1 | 1.3 | 5.9 | 168.2 | 8.5 |
| 10 - 20 cm | 5.6 | 0.9 | 3.6 | 188.5 | 7.8 |

**Table S3** OTU ID information. We accepted OTUs as belonging to Glomeromycota when they had a > 97.5 % similarity and > 99 % coverage than entries in MaarjAM.

| OTU ID | Accession | Description | Max score | Total score | Query coverage | E-value | Max identity |
| --- | --- | --- | --- | --- | --- | --- | --- |
| OTU5 | HF568031 | Glomeraceae Glomus Varela-Cervero15 BG9 | 154 | 218 | 90.10% | 5.20E-80 | 85.50% |
| OTU6 | HF568033 | Glomeraceae Glomus Varela-Cervero15 BG9 | 158 | 222 | 90.30% | 2.50E-82 | 84.70% |
| OTU11 | HF568087 | Glomeraceae Glomus Varela-Cervero15 BG21 | 135 | 200 | 89.00% | 5.10E-69 | 82.30% |
| OTU14 | LT723796 | Glomeraceae Glomus sp. VTX00342 | 34 | 34 | 15.70% | 1.40E-10 | 82.90% |
| OTU19 | JN009464 | Diversisporaceae Diversispora sp. VTX00380 | 516 | 516 | 100.00% | 0 | 99.40% |
| OTU22 | HF568033 | Glomeraceae Glomus Varela-Cervero15 BG9 | 158 | 222 | 90.30% | 2.50E-82 | 84.70% |
| OTU24 | KC665641 | Glomeraceae Glomus Lopez-Garcia14 Glo166 | 76 | 76 | 33.30% | 6.80E-35 | 83.80% |
| OTU26 | HF568087 | Glomeraceae Glomus Varela-Cervero15 BG21 | 173 | 238 | 90.10% | 5.20E-91 | 85.90% |
| OTU27 | LT723796 | Glomeraceae Glomus sp. VTX00342 | 34 | 34 | 15.70% | 1.40E-10 | 82.90% |
| OTU28 | KJ959950 | Glomeraceae Glomus sp. VTX00219 | 479 | 479 | 100.00% | 0 | 97.70% |
| OTU32 | HF568038 | Glomeraceae Glomus Varela-Cervero15 BG9 | 169 | 228 | 87.60% | 1.10E-88 | 85.40% |
| OTU34 | KC708365 | Claroideoglomeraceae Claroideoglomus sp. VTX00193 | 427 | 427 | 100.00% | 0 | 94.70% |
| OTU36 | FN869852 | Archaeosporaceae Archaeospora Aca VTX00338 | 473 | 473 | 100.00% | 0 | 98.80% |
| OTU37 | HF568087 | Glomeraceae Glomus Varela-Cervero15 BG21 | 149 | 216 | 91.50% | 4.10E-77 | 83.70% |
| OTU39 | LT934587 | Glomeraceae Glomus sp. | 158 | 225 | 93.90% | 2.50E-82 | 83.30% |
| OTU41 | HF568087 | Glomeraceae Glomus Varela-Cervero15 BG21 | 176 | 238 | 89.40% | 9.60E-93 | 85.80% |
| OTU42 | HF568058 | Glomeraceae Glomus Varela-Cervero15 BG13 | 159 | 221 | 92.60% | 6.60E-83 | 84.00% |
| OTU45 | HF568021 | Glomeraceae Glomus Varela-Cervero15 BG7 | 122 | 190 | 87.00% | 1.80E-61 | 81.50% |
| OTU47 | KC665641 | Glomeraceae Glomus Lopez-Garcia14 Glo166 | 82 | 82 | 33.30% | 2.40E-38 | 85.10% |
| OTU50 | HF568040 | Glomeraceae Glomus Varela-Cervero15 BG9 | 174 | 238 | 86.50% | 1.40E-91 | 87.70% |
| OTU52 | HF568040 | Glomeraceae Glomus Varela-Cervero15 BG9 | 183 | 247 | 86.50% | 8.70E-97 | 88.70% |
| OTU54 | KC665641 | Glomeraceae Glomus Lopez-Garcia14 Glo166 | 85 | 85 | 33.30% | 4.50E-40 | 85.80% |
| OTU55 | HF568087 | Glomeraceae Glomus Varela-Cervero15 BG21 | 165 | 230 | 88.80% | 2.20E-86 | 85.30% |
| OTU58 | HF568034 | Glomeraceae Glomus Varela-Cervero15 BG9 | 164 | 220 | 87.90% | 8.50E-86 | 84.80% |
| OTU59 | HF568038 | Glomeraceae Glomus Varela-Cervero15 BG9 | 152 | 152 | 73.90% | 7.40E-79 | 83.90% |
| OTU60 | FN429106 | Glomeraceae Glomus VeGlo13 VTX00153 | 495 | 495 | 100.00% | 0 | 98.30% |
| OTU62 | AB746997 | Claroideoglomeraceae Claroideoglomus Yoshimura13b Glo17 | 435 | 435 | 100.00% | 0 | 94.30% |
| OTU63 | JQ246044 | Glomeraceae Glomus sp. VTX00359 | 79 | 79 | 67.60% | 1.30E-36 | 76.70% |
| OTU64 | LT934587 | Glomeraceae Glomus sp. | 152 | 214 | 92.10% | 7.50E-79 | 82.70% |
| OTU65 | LN900648 | Glomeraceae Glomus sp. VTX00096 | 477 | 477 | 100.00% | 0 | 97.30% |
| OTU66 | HF568040 | Glomeraceae Glomus Varela-Cervero15 BG9 | 166 | 227 | 87.00% | 6.00E-87 | 87.50% |
| OTU67 | AJ854097 | Archaeosporaceae Archaeospora sp. VTX00006 | 450 | 487 | 100.00% | 0 | 99.30% |
| OTU70 | EU417640 | Glomeraceae Glomus Sciaphila ledermannii symbiont VTX00166 | 498 | 498 | 100.00% | 0 | 98.50% |
| OTU71 | LN623488 | Paraglomeraceae Paraglomus MO-P2 VTX00433 | 125 | 195 | 87.60% | 3.20E-63 | 81.40% |
| OTU72 | LT983627 | Glomeraceae Glomus SS-G1 VTX00448 | 248 | 313 | 91.20% | 2.00E-134 | 92.30% |
| OTU73 | LN827079 | Glomeraceae Glomus sp. VTX00344 | 466 | 466 | 100.00% | 0 | 96.90% |
| OTU74 | HF568033 | Glomeraceae Glomus Varela-Cervero15 BG9 | 160 | 221 | 90.10% | 1.70E-83 | 84.90% |
| OTU75 | HF569099 | Gigasporaceae Scutellospora Varela-Cervero15 BS2 | 140 | 205 | 91.50% | 6.50E-72 | 82.30% |

**Appendix I** Defining stochasticity

The term stochasticity has often been used in the literature synonymously with variability [3]. Shoemaker *et al.* [3] propose narrowing down the definition of stochasticity to processes that can be represented in a probabilistic way defined by their parameters (e.g. mean, variance, and skew) and discriminate across three forms of stochasticity: demographic stochasticity, environmental stochasticity, and measurement error. Based on this definition it is possible that the aggregate of the community variance that is explained by deterministic processes and stochasticity falls below 100%.

In our study we define stochasticity in relation to the proportion of variance that is not explained by deterministic processes (i.e. deterministic processes and variance explain together a 100% of community variance). We do so because arbuscular mycorrhizal associations represent complex systems which we do not understand sufficiently to effectively model processes such as demographic stochasticity (priority effects and succession, for example, have been poorly defined for AMF systems) which could have led us to seriously underestimate stochasticity. We thus describe stochasticity here as the fraction of community variance that is not explained by deterministic processes [4].

To show a few examples of the proportion of stochasticity that authors often find in AMF communities we carried out a literature search. We used the key words "deterministic" AND "arbuscular" in the Web of Knowledge on 30 June 2020. The search yielded 9 results which we screened for studies that contained figures on variance explained following a variance partitioning exercise. There were four such studies, which we list below:

| study | variance explained |
| --- | --- |
| Maciel Rabelo Pereira *et al.* 2020 Journal of Biogeography, [5] | 9% |
| Rasmussen *et al.* 2018 New Phytologist [6] (Caruso 2018 New Phytologist Commentary, [7]) | 7-25% |
| Kohout *et al.* 2015 Molecular Ecology [8] | 39.3% |
| Dumbrell *et al.* 2010 ISME Journal [9] | 68.5% |

**Appendix II** Limitations of existing multivariate techniques in addressing spatio-temporal designs and how we addressed them

Most studies addressing the relative role of deterministic and stochastic processes in structuring communities engage into variance partitioning exercises with the aim to explain as much variance as possible (e.g. [10]). Accuracy of such estimates depends strongly on the degree to which the underlying ordination model captures the experimental design. For example, assaying a plot at multiple instances in time represents a violation of independence and can be potentially addressed through a repeated-measures design in which case plot will be the unit of the analysis (i.e. *Subject*) whereas time the within-subject factor (e.g. [11]). Determining between-subject and within-subject sources of variance is critical in the case of a variance partitioning exercise, because through this step it will become apparent that each group will be compared against a unique fraction of unexplained variance (e.g. within-subjects unexplained variance). Many of the studies addressing spatio-temporal designs, however, incorrectly assume that iterative harvests of a single plot are independent of each other.

Even though there are several techniques to address spatial autocorrelation in ordination analyses, to the best of our knowledge the only multivariate technique that works for temporal constraints is that of Palmer *et al.* [11], which was specifically proposed for split plot designs. To minimize the assumptions of our analyses we plot the data with a PCA (i.e. meaning that we do not propose any underlying model; Fig. 2a) and then calculate distributions of pairwise Bray-Curtis distances (Fig. 2b). To back up our analysis we fit a redundancy analysis model (Appendix III) in which we address temporal constraints by restricting permutations (and thus calculation of resulting *P* values) within plots. This approach may be an improvement compared to assuming full independence. For this reason, we engage into no variance partitioning exercise.

Our study was subject to some limitations which we would like to raise here. First, we did not assay soil properties and assumed that the abiotic parameters throughout the site were rather homogenous. To this end we included in our analysis spatial corrections which should have accounted for spatial autocorrelations in soil properties. Second to simultaneously address all factors we sacrificed the number of levels we considered. We worked with two plant species over two years and two seasons. Some of the results may be idiosyncratic on the choice we made. Third, we only assayed two AM plant species, which were the ones that showed sufficient abundance across the target forest site. One of them (*Euonymus)* could only be collected in the last harvest. This was done because there were only a few individuals of *Euonymus* in the forest site and their destructive harvest could modify meta-community dynamics of AMF species. As a result, to a certain degree the variance that we allocated to plant species was nested within the variance explained by season and year. To address this point, we only tested spatio-temporal parameters for *Hedera* and in the cases where we included *Euonymus* in analyses we formulated the models we fitted so that there were several alternative ranks of predictors.

**Appendix III** Statistical analyses

Most of our statistics were descriptive. Ordination analyses were carried out after Hellinger transforming community data to avoid distortions of distances in the Euclidean space (for example Fig. 2a). To assay the distributions of Bray Curtis distance we randomly paired samples 9999 times (Fig. 2b). Relative abundance figures of AMF families were descriptive (Fig. 2c).

We carried out an Analysis of Variance on the RDA models that we fitted. The default settings of the command anova.cca in R test hierarchically the terms of the RDA/CCA models and yield different results from when the predictors are fitted “simultaneously”. Additionally, the command does not offer possibilities to fully distinguish between within-factors and between factors variability (i.e. address repeated measures designs). An alternative is to constrain permutations so that they occur within the subjects of the analysis (i.e. here the plots), which we applied here. To address the fact that our experimental design was not balanced and that predictors were fitted hierarchically we tested alternative formulations of the models after changing the rank of the predictors. We additionally included models with and without the predictor *plant species*. To address spatial constraints, we used the Principal Coordinates of Neighbourhood Matrix (PCNM) approach to summarize space into three spatial axes. The parameters *year*, spatial PCNM axes and *plant species* (in the cases when it was included) were significant irrespective of their rank in the models. Season was never significant whereas the predictor *AM coverage* was only occasionally significant. We attach specific results:

**Redundancy Analysis Statistics**

```
> anova(myrda, by="terms", permutations=9999, strata=as.factor(field$plot))
Permutation test for rda under reduced model
Terms added sequentially (first to last)
Blocks:  strata
Permutation: free
Number of permutations: 9999

Model: rda(formula = data2 ~ AM_coverage + year + season + plant_species + PCNM1 + PCNM2 + PCNM3, data = field, scale = T)
          	Df Variance  	F Pr(>F)
AM_coverage	1   0.6071 1.5900 0.0002 ***
year 	      1   0.8247 2.1600 0.0009 ***
season     	1   0.5360 1.4040 0.1974
plant_species  1   0.5651 1.4801 0.0203 *
PCNM1      	1   0.5948 1.5580 0.1529
PCNM2      	1   0.5199 1.3616 0.5799
PCNM3      	1   0.5720 1.4982 0.0351 *
Residual  	78  29.7804
---
Signif. codes:  0 ‘***’ 0.001 ‘**’ 0.01 ‘*’ 0.05 ‘.’ 0.1 ‘ ’ 1

> anova(myrda, by="terms", permutations=9999, strata=as.factor(field$plot))
Permutation test for rda under reduced model
Terms added sequentially (first to last)
Blocks:  strata
Permutation: free
Number of permutations: 9999

Model: rda(formula = data2 ~ year + plant_species + AM_coverage + season + PCNM1 + PCNM2 + PCNM3, data = field, scale = T)
          	Df Variance  	F Pr(>F)
year       	1   0.7991 2.0929 0.0011 **
plant_species  1   0.6027 1.5786 0.0132 *
AM_coverage	1   0.6111 1.6005 0.1137
season     	1   0.5200 1.3619 0.2276
PCNM1      	1   0.5948 1.5580 0.1459
PCNM2      	1   0.5199 1.3616 0.5852
PCNM3      	1   0.5720 1.4982 0.0344 *
Residual  	78  29.7804
---
Signif. codes:  0 ‘***’ 0.001 ‘**’ 0.01 ‘*’ 0.05 ‘.’ 0.1 ‘ ’ 1

> myrda<-rda(data2~year+season+AM_coverage+PCNM1+PCNM2+PCNM3, data=field[1:71,], scale=T)
> anova(myrda, by="terms", permutations=9999, strata=as.factor(field$subplot)[1:71])
Permutation test for rda under reduced model
Terms added sequentially (first to last)
Blocks:  strata
Permutation: free
Number of permutations: 9999

Model: rda(formula = data2 ~ year + season + AM_coverage + PCNM1 + PCNM2 + PCNM3, data = field[1:71, ], scale = T)
        	Df Variance  	F Pr(>F)
year     	1   0.7230 1.5365 0.0364 *
season   	1   0.5508 1.1706 0.3986
AM_coverage  1   0.7495 1.5929 0.0459 *
PCNM1    	1   0.7039 1.4961 0.6544
PCNM2    	1   0.4805 1.0212 0.6495
PCNM3    	1   0.6796 1.4445 0.0937 .
Residual	64  30.1127
---
Signif. codes:  0 ‘***’ 0.001 ‘**’ 0.01 ‘*’ 0.05 ‘.’ 0.1 ‘ ’ 1

> anova(myrda, by="terms", permutations=9999, strata=as.factor(field$plot)[1:71])
Permutation test for rda under reduced model
Terms added sequentially (first to last)
Blocks:  strata
Permutation: free
Number of permutations: 9999

Model: rda(formula = data2 ~ year + season + PCNM1 + PCNM2 + PCNM3 + AM_coverage, data = field[1:71, ], scale = T)
        	Df Variance  	F Pr(>F)
year     	1   0.7230 1.5365 0.0338 *
season   	1   0.5508 1.1706 0.4004
PCNM1    	1   0.7231 1.5369 0.7505
PCNM2    	1   0.4844 1.0294 0.1324
PCNM3    	1   0.9037 1.9206 0.0587 .
AM_coverage  1   0.5023 1.0676 0.9381
Residual	64  30.1127
---
Signif. codes:  0 ‘***’ 0.001 ‘**’ 0.01 ‘*’ 0.05 ‘.’ 0.1 ‘ ’ 1

> anova(myrda, by="terms", permutations=9999, strata=as.factor(field$subplot)[1:71])
Permutation test for rda under reduced model
Terms added sequentially (first to last)
Blocks:  strata
Permutation: free
Number of permutations: 9999

Model: rda(formula = data2 ~ PCNM1 + PCNM2 + PCNM3 + AM_coverage + season + year, data = field[1:71, ], scale = T)
        	Df Variance  	F Pr(>F)
PCNM1    	1   0.7513 1.5968 0.0788 .
PCNM2    	1   0.4173 0.8869 0.0788 .
PCNM3    	1   0.8174 1.7372 0.0788 .
AM_coverage  1   0.5426 1.1533 0.0788 .
season   	1   0.5270 1.1201 0.4780
year     	1   0.8316 1.7674 0.0321 *
Residual	64  30.1127
---
Signif. codes:  0 ‘***’ 0.001 ‘**’ 0.01 ‘*’ 0.05 ‘.’ 0.1 ‘ ’ 1
```

**Repeated-measures ANOVA**

```
> summary(aov(S~AM_coverage+as.factor(plant_species)+as.factor(season)+as.factor(year), Error=(as.factor(subplot)/as.factor(sampling)), data=field))
                         Df Sum Sq Mean Sq F value   Pr(>F)
AM_coverage               1    0.3    0.33   0.048 0.827609
as.factor(plant_species)  1   80.9   80.92  11.701 0.000981 ***
as.factor(season)         1    3.1    3.07   0.443 0.507397
as.factor(year)           1    1.3    1.34   0.194 0.660641
Residuals                81  560.2    6.92
---
Signif. codes:  0 ‘***’ 0.001 ‘**’ 0.01 ‘*’ 0.05 ‘.’ 0.1 ‘ ’ 1
```

**Indicator Species Analysis**

```
> ind.sp = multipatt(data2, as.factor(paste0(as.factor(field$plant_species), as.factor(field$AM_coverage))), control = how(nperm=9999), max.order=2, restcomb=c(1:6, 9:10))

 Association function: IndVal.g
 Significance level (alpha): 0.05

 Total number of species: 34
 Selected number of species: 5
 Number of species associated to 1 group: 2
 Number of species associated to 2 groups: 3
 Number of species associated to 3 groups: 0

 List of species associated to each combination:

 Group Euonymus europaea  #sps.  1
       stat p.value
OTU70 0.826   2e-04 ***

 Group Hedera helix  #sps.  1
       stat p.value
OTU19 0.644  0.0383 *

 Group Euonymus europaea+Euonymus europaea  #sps.  1
      stat p.value
OTU2 0.854  0.0349 *

 Group Hedera helix+Hedera helix  #sps.  2
       stat p.value
OTU8  0.904  0.0007 ***
OTU13 0.880  0.0027 **
---
Signif. codes:  0 ‘***’ 0.001 ‘**’ 0.01 ‘*’ 0.05 ‘.’ 0.1 ‘ ’ 1
```
